# Testing the parasite mass burden effect on host behaviour alteration in the *Schistocephalus*-stickleback system

**DOI:** 10.1101/222802

**Authors:** Lucie Grécias, Julie Valentin, Nadia Aubin-Horth

## Abstract

Many parasites with complex life cycles modify their intermediate host’s behaviour, which has been proposed to increase transmission to their definitive host. This behavioural change could result from the parasite actively manipulating its host, but could also be explained by a mechanical effect, where the parasite’s physical presence affects host behaviour. We created an artificial internal parasite using silicone injections in the body cavity to test this *mechanical effect* hypothesis. We used the *Schistocephalus solidus -* threespine stickleback (*Gasterosteus aculeatus*) system, as this cestode can reach up to 92% of its fish host mass. Our results suggest that the mass burden brought by this macroparasite alone is not sufficient to cause behavioural changes in its host. Furthermore, our results show that wall-hugging (thigmotaxis), a measure of anxiety in vertebrates, is significantly reduced in *Schistocephalus*-infected sticklebacks, unveiling a new altered component of behaviour that may result from manipulation by this macroparasite.

## Introduction

Many parasites have complex life cycles with multiple hosts, and alter various aspects of their host’s biology (Poulin, 2010). Infection can be accompanied by changes in morphology, physiology, life history and behaviour of the host (Poulin and Thomas, 1999). Those phenotypic modifications can stem from non-exclusive causes: specific manipulation, side effects (pathological consequences) or by-products, which can be coincidentally beneficial (Thomas et al., 2005). Understanding these mechanisms will shed light on host–parasite interactions and coevolution (Poulin, 2007). However, in most host-parasite systems, the proposed causes of those changes are not tested using experimental confirmation (Moore, 1983). This is in part because it is technically complicated to do so, as multiple proximal causes may intervene to result in these altered host phenotypes (Grécias et al., 2017).

In host-parasite systems where the parasite infects its host internally, the presence of this foreign body could cause modification in the host phenotype, including its behaviour. The parasite’s “mechanical effect” can be defined as all effects induced merely by the physical presence of the parasite or its movements that have consequences on host phenotype in the time window in which they are exerted by the parasite. For example, the metacercariae of the *Tylodelphys* trematode infect the vitreous humour of the eyes of its fish host (Stumbo and Poulin, 2016). The parasite is not encysted and can cross from one side of the eye to the other. This trematode shows a daily rhythm of movement within the eye, obstructing its host’s retina during the day when its final fish-eating bird host is active, while leaving the eye fully functional at dusk and night. This example shows the combined effect of mechanical and parasite behaviour as a cause of potential host behaviour modification. Another example of manipulative parasite that might change its host behaviour through a mechanical modification combined with its behaviour is the trematode *Leucochloridium paradoxum*, which forms broodsacs that migrate to the tentacles of terrestrial snails and pulsate rapidly. Thanks to this parasite presence and behaviour, the snails’ eye stalks resemble caterpillars, resulting in increased predation by the definitive bird host (Wesołowska and Wesołowski, 2014). In both cases, the parasite’s behaviour (circadian movement in the eye, migrating to eye stalk and pulsating) is coupled with the effect of physical presence to affect the host’s phenotype, but it is conceivable that the sole parasite presence could affect its host.

In some cases, the mass of the internal parasite approaches the mass of its host (Arme and Owen, 1967). The threespine stickleback (*Gasterosteus aculeatus*) can be infected by a tapeworm, *Schistocephalus solidus.* This macro parasite can attain 40% of the host weight and, as multiple infections are common, combined worms can weigh up to 92% of the host weight (Hopkins and Smyth, 1951). The space taken by the parasite in the body cavity of the host could have major impact on the fish physiology. Growth of visceral organs is restricted because of the lack of space (Pascoe and Mattey, 1977; Walkey and Meakins, 1970). The stomach’s capacity for expansion may thus be limited in heavy infections (Milinski, 1985), potentially resulting in a decrease in food intake (Wright et al., 2006).The fish’s abdomen becomes grossly distorted as the parasite grows inside it (Arme and Owen, 1967). This abdomen distortion may make sticklebacks more vulnerable to predatory birds that select the largest available fish from the population (Van der Veer et al., 1997). *S. solidus* might also alter the buoyancy of its fish host, as the density of the cestode is higher than that of the stickleback (LoBue and Bell, 1993). This means that the host must use its swim bladder to reach higher buoyancy despite the parasite mass (Talarico et al., 2017). Concurrently with morphological changes, a suite of behaviours is changed in sticklebacks harbouring at least one infective parasite. They show more risky behaviours: they lose their anti-predator response and forage at a higher rate, even under the risk of predation (Milinski, 1985; Giles, 1987; Godin and Sproul, 1988; Barber et al., 2004) (Tierney et al., 1993). They also spend less time swimming with a group than uninfected ones when satiated (Barber et al., 1995; Barber et al., 1998). Moreover, infected fish tend to stay at the water surface both in nature and in the laboratory, contrary to healthy sticklebacks (they show “reversed geotaxis”, LoBue and Bell, 1993; Quinn et al., 2012). Studies on host manipulation have aimed to unravel the most plausible cause of host behaviour modification through manipulation of the neuroendocrine state of infected stickleback (Grécias et al. 2017) or by directly measuring monoamine levels in parasitized sticklebacks compared to healthy ones (Øverli et al., 2001). Bioinformatics and genomics studies have also shown that certain *S. solidus* proteins (such as Wnt4, known for its role in signal transduction in other systems) proposed to be a mimic of the stickleback protein, are expressed at the infective stage in *S. solidus* (Hébert et al., 2015; Hébert et al., 2016). However, to our knowledge, no study has tested the mechanical effect hypothesis of host alteration by a parasite using an experimental approach, in this system or another.

The aim of this study was to assess the impact of a parasite mass burden on the behaviour of its host in order to test whether there is a causal link between the parasite “mechanical effect” and behaviour alteration in the stickleback-cestode system. We used phenotypic engineering, the experimental modification of an organismic phenotype, as it allows to separate a complex phenotype into its parts and to focus on the effects of a single trait (Kliman, 2016) that allows testing predictions in a specific manner (Andersson, 1992; Aubret et al., 2003). We manipulated the space available in the body cavity of the fish by injecting an artificial silicone “worm” that is inert but similar in physical property to a live parasite, allowing isolating the mechanical effect of *Schistocephalus* on its host. We quantified four different behaviours: reversed geotaxis (tendency to use the top of the water column), response to a predator, feeding, and thigmotaxis (wall-hugging tendency). We predicted that sticklebacks harbouring an artificial worm would increase the time they spend swimming at the surface (reversed geotaxis) because of their buoyancy, as infected sticklebacks do. We also predicted that the sticklebacks harbouring an artificial worm would decrease their latency to start feeding, to freeze after an attack (they would freeze immediately instead of displaying a typical fleeing anti-predator response, because of an impairment in locomotion) and to resume moving. Finally, we measured a behaviour not previously quantified in infected sticklebacks, the tendency to avoid exposure to potential threats by hugging walls in an open field (thigmotaxis or centrophobic behaviour). This behaviour is used as a measure of anxiety in mammals, fish and insects (Maximino et al., 2010; Webster and Laland, 2015; Mohammad et al., 2016) and has been observed in other infected hosts (Poulin, 2001). We predicted that infected sticklebacks would have a low thigmotaxis (they would spend less time near the walls of the aquarium and more time in the center) and that sticklebacks harbouring artificial worms would have the same behaviour.

## Materials and Methods

### Sampling

We caught *Gasterosteus aculeatus* juveniles (n=40) from Lac Témiscouata (47°40′33″N 68°50′15″O, Québec, Canada, August 2015) and kept them for 10 months in a common environment before the behavioural tests. They were kept with the same conditions (1-2 fish per 2 liters) under a natural lightdark cycle mimicking their natural environment (Québec City, Québec, Canada) and a water temperature of 15 °C. Fish were fed every morning with a mix of blood worms and artemia.

### Experimental treatments

To evaluate the influence of parasite presence on fish behaviour, we implanted an artificial worm in uninfected sticklebacks. We used an injection of silicone (Mold Star 16 – Fast, Smooth-On, Inc., USA) to simulate the presence of the parasite in the body cavity of its host. The use of silicone is non-toxic and common in endocrinology studies, see example in Ros et al., 2012). This specific type of silicone was selected for its similarity in density with *S. solidus* (see Fig. S1 for a direct comparison). To replicate the mass of the parasite, we injected 80 ul of silicone per injection every two days on three occasions (see below) to achieve a 150 mg mass. In parallel, to discriminate the effect of the mere exposure to silicone on behaviour from the effect of the volume of the artificial worm, we injected a small amount of silicone to a separate group of fish used as controls (15 ul / injection, for a total mass of 30 mg), also every two days on three occasions.

We had four experimental treatments: Silicone Artificial Parasite (80 ul of silicone / injection, n=15), Silicone Control (15 ul of silicone / injection, n=14), infected fish (infected with *S. solidus*, n=5) and control fish (no material injected, n=6). Infected and control fish were handled in the same way as other groups, but instead of injecting them, we only inserted a needle in their body cavity.

Fish were isolated in a 2L tank at the start of the experiment (Day 1). We assigned them randomly to a treatment for the rest of the experiment. Fish were fed every day, except on the day before the behavioural tests. Fish were handled every two days. During the first week (Control week) we handled the fish to recreate the experimental manipulation, but without any injection (day 1, 3, and 5). Fish were taken out of the tank, held in a moist sponge on their back to have easy access to the body cavity. At the end of the Control week (Day 7), we measured their behaviour (see “behavioural experiments” below) and placed them back in their tank where they were fed. The second week (Engineering week) started on day 8. Handling procedure was the same, but we injected the fish according to their treatment (day 8, 10 and 12). At the end of the Engineering week (Day 14), we measured the same behaviours. On day 15, we euthanized the fish. Total length (6.6 ± 0.59 cm), mass (1.6 ± 0.84 g), sex, and the presence of a parasite were noted (a single infective plerocercoid (>50mg) was found inside the body cavity of each infected stickleback)

### Behavioural experiments

Two cameras (on the top and side of the aquarium) recorded all behavioural trials in the test aquarium (30Wx30Dx90L cm, 20cm of water depth).

At the start of the experiment, an isolated focal fish was placed inside an opaque container (500 mL), which was placed horizontally into the behaviour test tank. As soon as the fish exited the container (or after 150 sec), the container was taken out of the aquarium and the behavioural trial started. The first 30 seconds after the container removal were not used because of the water movement due to its removal.

Thigmotaxis: Latency to enter the center of the tank and time spent in the center of the tank (at more than 5 cm of the edge of the aquarium) were analyzed during 120 seconds.

Geotaxis: The time spent near the surface (in the 10 cm upper zone) was also quantified during this 120-second period.

Latency to feed: Right after the preceding measurements, the fish was fed with blood worms released in the water using a plastic pipette (within 5 cm of the fish head). Food was released in front of the fish to ensure it was detected in all trials. Latency to feed was quantified, up to a maximum of 150 seconds.

Response to a predator: As soon as the fish had fed (or after 150 sec), we attacked the fish (within 5 cm of the fish head) with an artificial heron head (only the beak penetrated the water, 10 to 15 cm) to simulate a predator attack. We quantified the time taken to freeze after the attack, and the time spent frozen until activity was resumed, up to 150 seconds.

### Behavioural and statistical analysis

Latency to enter the center and time spent in the center, as well as time spent near the surface were analyzed using an automatic tracking module in the Ethovision software (Ethovision XT 11.5, Noldus Information Technology, Wageningen, The Netherlands, Noldus et al., 2001). Latency to feed, latency to freeze after a predator attack and latency to resume activity after the attack were analyzed by hand. All analyses were done while blind to treatment.

We performed all statistical analyses in R software version 3.0.2 (R Foundation for Statistical Computing, Vienna, Austria). We conducted analyses of variance on aligned rank transformed data with the package “ARTool” (Wobbrock et al., 2011). We aligned and rank transformed our data and ran a nonparametric ANOVA. Individuals, weeks and treatments and their interaction (treatment*week) were added to the statistical model. To account for variation in behaviour between individual sticklebacks even before any manipulation, we included tank, mass, length and sex of the fish along with the parasite weight as fixed effects, and fish id as a random effect (for repeated measurements). We checked the effect of tank, mass, length, sex, parasite weight with visual validation of multiple PCAs and no effects (clusters) were found on any behaviour. We then excluded those factors from the model. Finally, we did a post hoc pairwise comparison with the package “lsmeansLT” (Lenth, 2016) which reports p-values that are Tukey-corrected for multiple comparisons. We report significant differences in these post-hoc tests.

## Results and discussion

Infection-associated changes in host behaviour are often explained as direct manipulation by the parasite that facilitates life cycle completion, often by increasing transmission rates to the final host. However, there are alternative explanations for these observations that do not necessarily invoke any active manipulation. We designed the experiment to test the impact of the parasite mass burden on stickleback behaviour and to compare it to the effect of a *S. solidus* infection using an artificial worm. Contrary to our predictions, phenotypically manipulated individuals with an artificial worm mass did not show similar behavioural responses to infected ones. In fact, the largest difference we quantified between treatments was between infected fish and fish injected with an artificial worm. Infected fish spent more time performing risky behaviour while fish implanted with an artificial worm did not differ from control fish. Thigmotaxis was different between infected and phenotypically-engineered fish: infected fish spent significantly more time in the center than fish with an artificial parasite (p = 0.007) and fish used a silicone controls (p = 0.039) (Fig. 1a, Table S1 and S2). Infected fish also had a shorter latency to enter the center of the test tank than the artificial parasite-implanted fish, but this difference was not significant (post-hoc test, p = 0.055) (Fig. 1b, Table S1 and S2). Geotaxis was not affected: there were no differences in the time spent in the upper part of the aquarium across any of the treatments (Fig. S2, Table S1 and S2). Feeding also differed between groups: infected fish had a significantly shorter latency to feed than fish with the artificial worm (p = 0.022) and fish used as silicone controls (p = 0.031) (Fig. 1c, Table S1 and S2). Furthermore, infected fish had a shorter latency to freeze after a predator attack than the artificial parasite-implanted fish (p = 0.0003) and the fish injected with a small amount of silicone (p = 0.008) (Fig. 1d, Table S1 and S2). There were no significant differences in the time spent frozen after an attack across any of the treatments (Fig. S3, Table S1). Our results show that the experimental behavioural tests were appropriate to detect previously described changes in behaviour in infected fish, but that the phenotypic engineering approach using silicone injections did not result in the predicted behavioural changes. Our results also show that thigmotaxis (wall hugging, a measure of anxiety in several vertebrates) is significantly lowered in infected sticklebacks, a behaviour change not previously reported for this system that can be reliably quantified and used in future studies.

**Figure 1.**
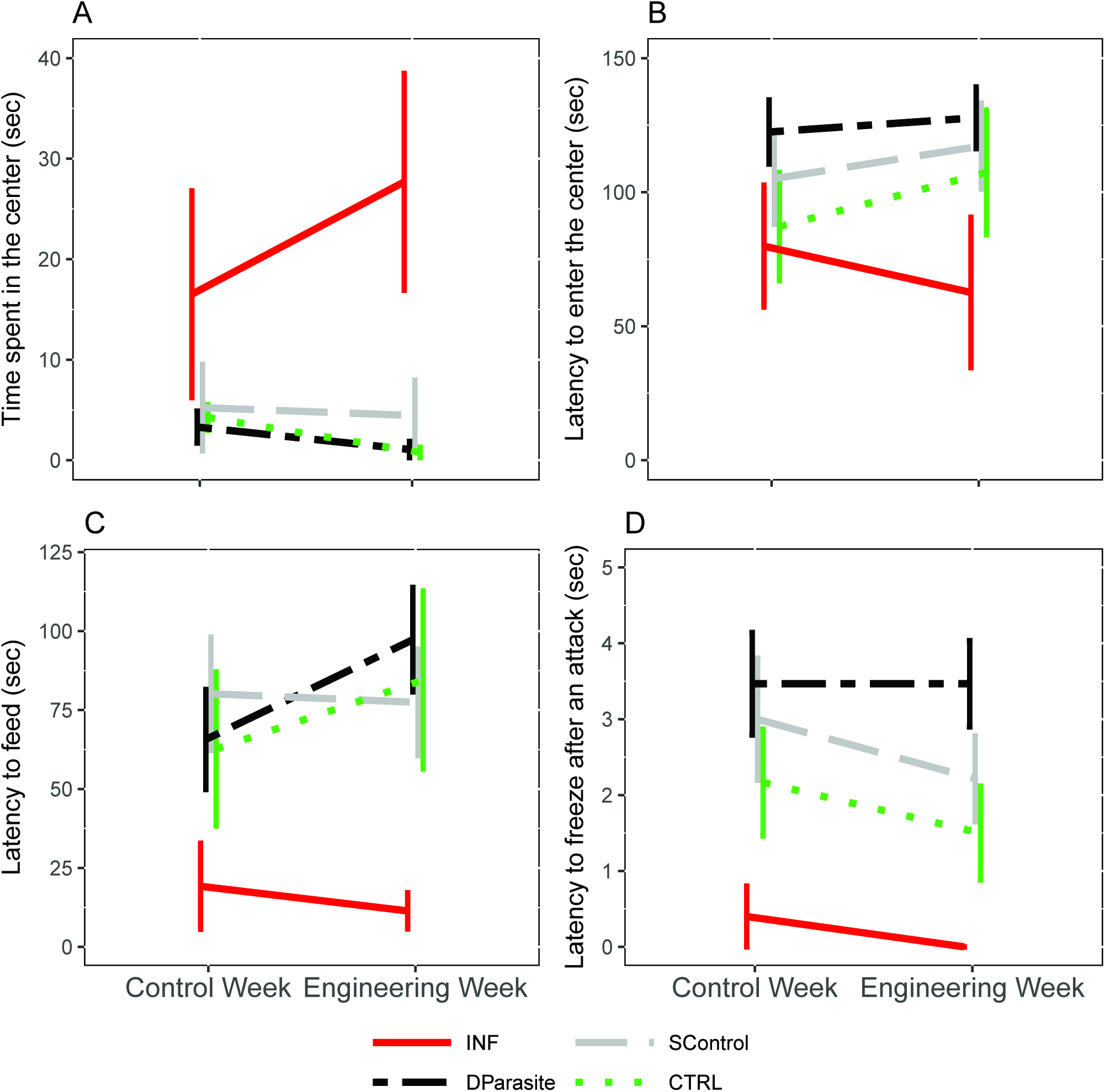
Phenotypic engineering with an artificial worm does not recreate the behaviour of a *S. solidus* infected fish. Infected fish spent more time performing risky behaviours while fish implanted with an artificial worm did not differ from control fish. Average behavioural response and standard error are presented for each experimental treatment (artificial parasite, infected fish, silicone control, and control fish) and week of treatment (control week and engineering week). (a) Thigmotaxis, time spent in the center of the tank, (b) Latency to enter the center of the tank, (c) Latency to feed, and (d) Latency to freeze after an attack. DP: dummy parasite (dash-dotted black line, n=15), INF: infected (solid red line, n=5), SC: silicone control (dashed grey line, n=14), and CTRL: non-infected (dotted green line, n=6).

This study was the first to test a strictly mechanical effect hypothesis to explain behavioural alteration in a parasitized host. Our results suggest that the space taken by a *Schistocephalus* worm at its infective stage when behavioural modifications are usually measured is not the cause of the behavioural changes in its host, at least not in isolation. Indeed, *S. solidus* is alive in the body cavity of its host, and hence is not only a mass burden but is also moving and can impact fish physiology, including the digestive track, when moving between organs. We did not try to reproduce the movement of the parasite in the body cavity of its host as a cause of behavioural modification. Therefore, the “parasite behaviour” component of the mechanical effect hypothesis remains to be tested alone and in combination with other causes such as the immune response, hunger or neuromodulation (Grécias et al., 2017). Even though we did not measure significant differences in behaviour between the two volumes of silicone treatments used, we suggest that the treatment where a small amount of material is injected is an essential control to have to include in phenotypic engineering studies trying to mimic the mass of a parasite with inert substances.

Sticklebacks captured during the day near the surface of a lake have a greater likelihood of being infected with *S. solidus* and have larger parasites than infected fish caught at the surface only at night (Quinn et al., 2012). Furthermore, in the laboratory, infected sticklebacks spend significantly more time foraging in the upper part of an experimental tank than control fish (Talarico et al., 2017). However, contrary to our prediction, we did not find significant differences in time spent at the surface between groups (Fig. S2). In a previous laboratory experiment, we tested if individuals prefer to swim in shallow waters versus in the deeper part of a tank and showed variation among treatments and also among individuals from a given treatment (Grécias et al., 2017). We expected that allowing individuals to use the full water column (rather than giving them a binary choice) would result in a better discrimination of geotaxis, but it did not in the experimental set-up used, which was much shallower than in Talarico et al. 2017 (20 cm water depth in the present study versus 50 cm). As a cautionary note, we only had five infected fish available for the experiment, which limited our statistical power to detect a geotaxis tendency if this behaviour is more variable among individuals. Furthermore, we did not use experimental infections but rather relied on wild infected-individuals. Another study showed no differences in the water column position at which parasitized and healthy sticklebacks were caught, and propose that buoyancy problems may increase with larger parasite mass, but that the worms do not usually attain this mass in the wild because its host is predated earlier (Ness and Foster, 1999). In freshwater fish, the swim bladder takes 7% of the body volume (Fänge, 1983). While it has been reported that there is no significant reduction in swim bladder size of infected fish (Arme and Owen, 1967), it would be interesting to not only analyze the space taken by the parasite as in the present study, but also its impact on the swim bladder, as previously suggested by Meakins and Walkey (1975).

The “wall hugging tendency” (thigmotaxis) was lower in infected sticklebacks, as they swam readily in the center of the tank, which could reflect lower anxiety brought-about directly or indirectly by the *Schistocephalus* infection. Thigmotaxis has been previously measured in wild healthy ninespine sticklebacks (*Pungitius pungitius*), a closely related species to threespine sticklebacks, and has been shown to be consistent when measured more than once on the same individual (Webster and Laland, 2015). Wall-hugging ninespine sticklebacks (high thigmotaxis) also showed lower activity and a higher latency to explore a new environment (Webster and Laland, 2015). In zebrafish, avoidance behaviour measured as thigmotaxis is used to assess anxiety level, and can be lowered using anxiety-lowering drugs or ethanol exposure (Maximino et al., 2010). The quantitative assay we used in this study could thus be used in future studies of the mechanistic proximal causes of behavioural alteration in *Schistocephalus-*infected sticklebacks. Furthermore, additional investigation of anxiety alteration in these infected individuals is warranted.

Modification of host behaviour by parasites is a fascinating but controversial phenomenon, since it is often difficult to disentangle if it is an adaptation of the parasite or simply a side-effect of the infection that has no fitness consequence for both the host and the parasite (Thomas et al., 2005). Our findings suggest that the use of silicone might not be the correct substitute to recreate the parasitic mass of *S. solidus.* We thus cannot conclude nor reject that the mechanical hypothesis is a possible cause of behaviour modification by a parasite. However, the general approach of using phenotypic engineering to create artificial parasites is a potentially fruitful avenue to dissect the proximal causes of parasitic manipulation.

## Ethics statement

All work was carried out in compliance with Animal Care and Use Guidelines, under a permit of the Comité de Protection des Animaux de l’Université Laval (CPAUL, permit 2014-069-3).

## Data accessibility

The complete dataset is available as supplementary material

## Competing interests

The authors declare no competing or financial interests.

## Authors’ contributions

L.G. designed the study with input from N.A.-H. L.G. performed the manipulation experiments, carried out the statistical analysis. L.G. and J.V. analyzed the behaviours. L.G. and N.A.-H. drafted the manuscript. All authors have reviewed and agreed on the final manuscript.

## Acknowledgments

We thank the LARSA employees for help with the fish rearing and Sergio Cortez Ghio for help with statistical analysis. We thank Katie Piechel, Jacques Brodeur, Christian Landry and Julie Turgeon for discussions that lead to this project. Chloé Berger and Verônica Alves provided comments on a previous version of this manuscript.

## Funding

This study was funded through a Fonds de Recherche du Québec - Nature et Technologies (FRQ-NT) Programme de Recherche en Équipe grant to N.A.-H. and a Natural Sciences and Engineering Research Council of Canada (CRSNG) grant through the Discovery grant program to N.A.-H.

## Supplementary tables

**Table S1.** Statistical parameters of ANOVAS testing the effect of treatment, week of measurement and their interaction on a given behaviour.

| Behaviour | Results ANOVA |  |  |  |
| --- | --- | --- | --- | --- |
|  | Factor | F | Df | p-value |
| Latency to enter the center | Treatment | 3.24 | 3 | 0.03 |
|  | Week | 1.59 | 1 | 0.21 |
|  | Treatment*week | 0.80 | 3 | 0.50 |
| Time in the center | Treatment | 5.24 | 3 | 0.0042 |
|  | Week | 1.22 | 1 | 0.28 |
|  | Treatment*week | 2.17 | 3 | 0.11 |
| Time spent near the surface | Treatment | 2.99 | 3 | 0.44 |
|  | Week | 0.41 | 1 | 0.53 |
|  | Treatment*week | 1.14 | 3 | 0.34 |
| Latency to feed | Treatment | 3.20 | 3 | 0.035 |
|  | Week | 0.60 | 1 | 0.44 |
|  | Treatment*week | 0.19 | 3 | 0.90 |
| Latency to freeze | Treatment | 6.49 | 3 | 0.001 |
|  | Week | 0.45 | 1 | 0.50 |
|  | Treatment*week | 0.14 | 3 | 0.93 |
| Time spent frozen | Treatment | 2.48 | 3 | 0.076 |
|  | Week | 3.86 | 1 | 0.057 |
|  | Treatment*week | 0.87 | 3 | 0.46 |

**Table S2.** Statistically significant comparisons between two treatments for a given behaviour test. INF: infected. DP: artificial worm. SC: silicone control.

| Behaviour | Results post hoc tests |  |  |  |
| --- | --- | --- | --- | --- |
|  | Comparisons | Estimate | Std. Error | p-value |
| Latency to enter the center | INF - DP | -22.3 | 8.62 | 0.0554 |
| Time in the center | INF – DP | 25.53 | 7.77 | 0.0084 |
|  | INF - SC | 28.38 | 7.84 | 0.003 |
| Time spent near the surface | INF– SC | -24.67 | 9.63 | 0.0601 |
| Latency to feed | INF– DP | -24.8 | 8.41 | 0.0218 |
|  | INF – SC | -23.9 | 8.48 | 0.0306 |
| Latency to freeze | INF– DP | -36.3 | 8.36 | 0.0003 |
|  | INF – SC | -27.8 | 8.43 | 0.0082 |

## Supplementary figures

**Figure S1.**
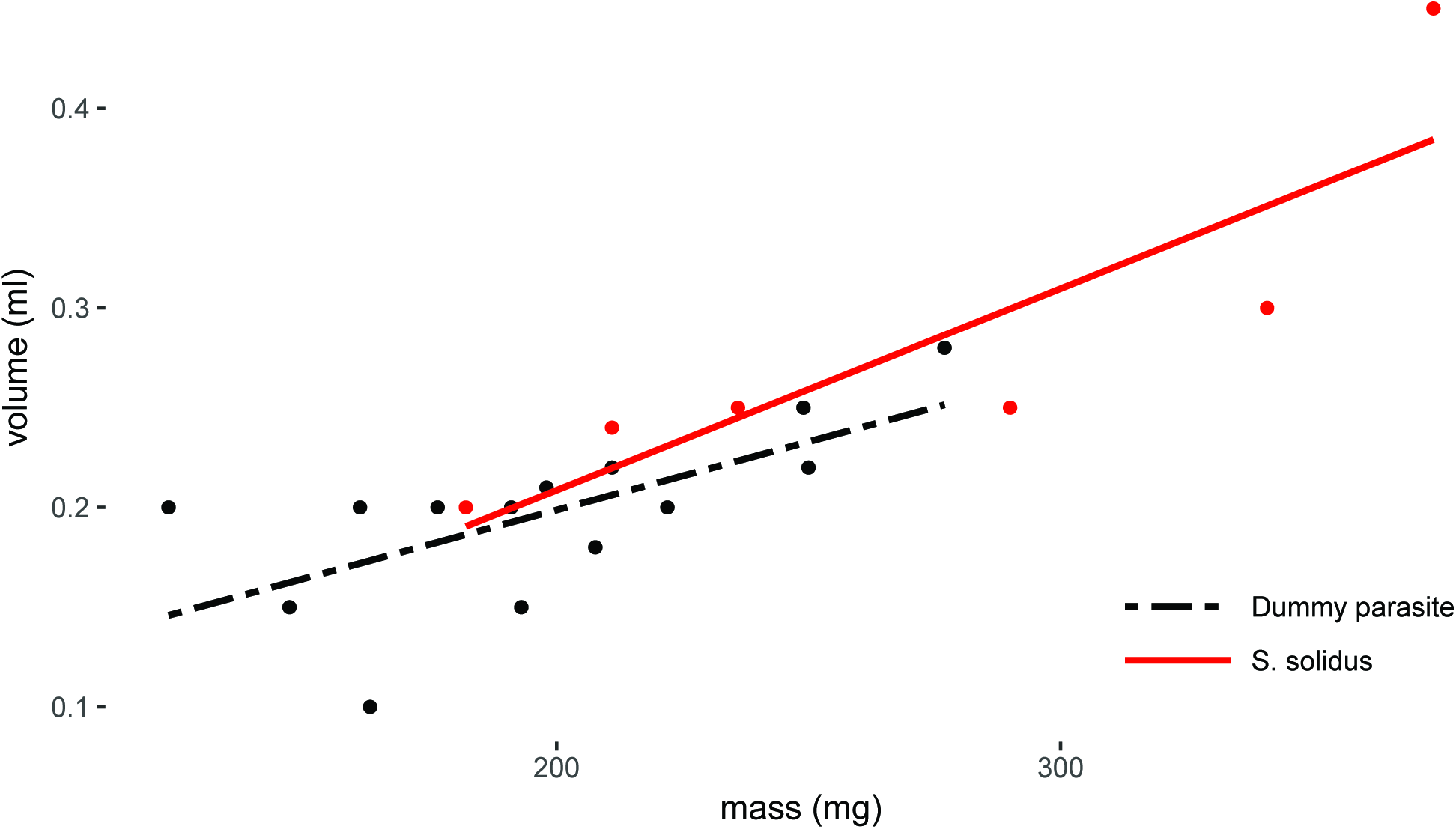
The artificial parasite and *S. solidus* worms have similar densities. Relationship between mass (mg) and volume (ml) of the silicone-made dummy parasite (dash-dotted black line) and the parasite *S. solidus* (solid red line).

**Figure S2.**
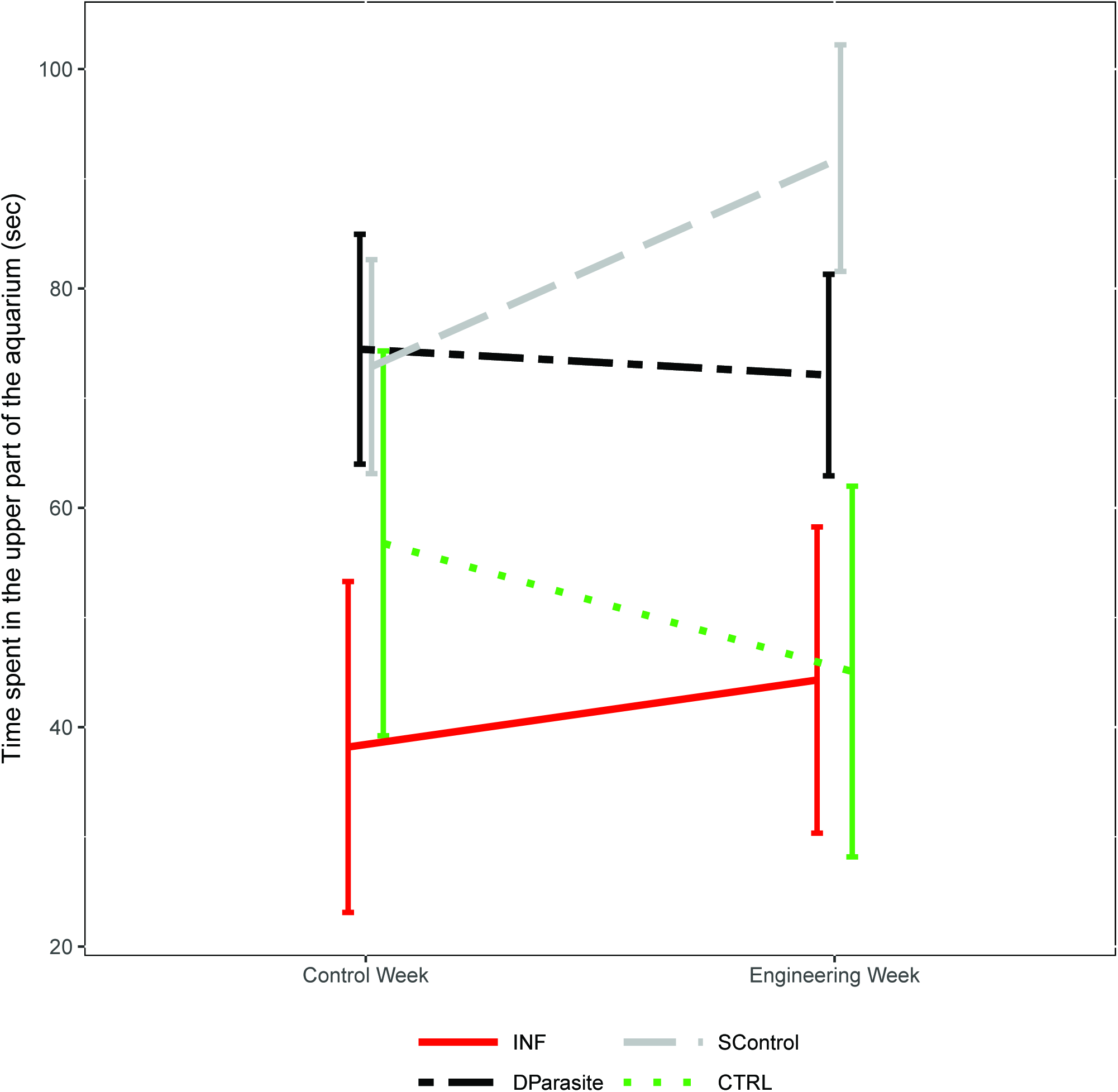
Geotaxis did not differ significantly between treatments. Time spent in the upper section of the aquarium (average and standard error) for each experimental treatment (artificial parasite, infected fish, silicone control, and control fish) and week of treatment (control week and engineering week). DP: dummy parasite (dash-dotted black line, n=15), INF: infected (solid red line, n=5), SC: silicone control (dashed grey line, n=14), and CTRL: non-infected (dotted green line, n=6).

**Figure S3.**
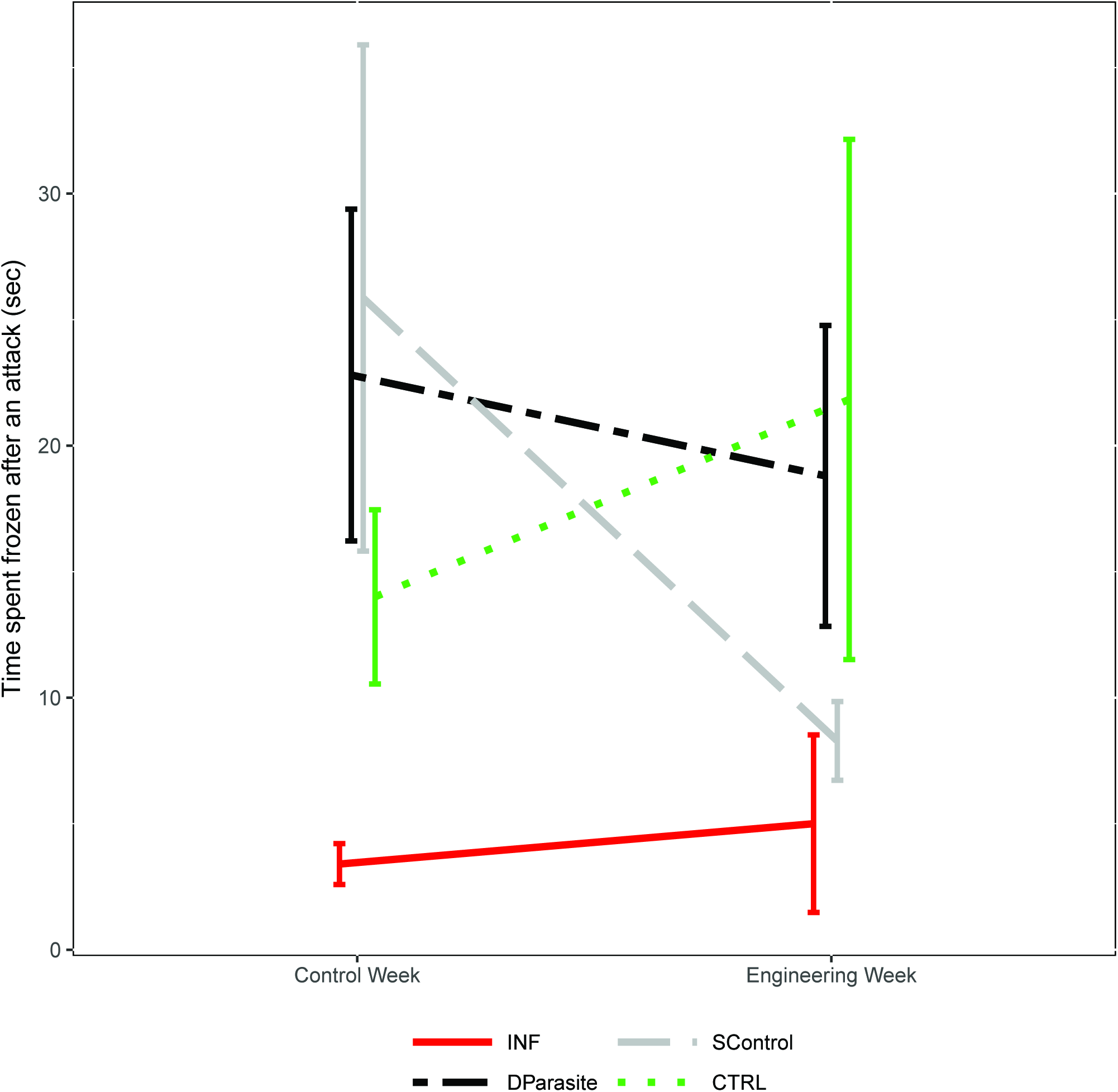
Freezing response to a predator presence did not differ significantly between treatments. Time spent frozen after an attack (average and standard error) for each experimental treatment (artificial parasite, infected fish, silicone control, and control fish) and week of treatment (control week and engineering week). DP: dummy parasite (dash-dotted black line, n=15), INF: infected (solid red line, n=5), SC: silicone control (dashed grey line, n=14), and CTRL: non-infected (dotted green line, n=6).

